# Multidisciplinary field research in Kabwe, Zambia, towards better understanding of lead contamination of the city - A short report from a field survey

**DOI:** 10.1101/096164

**Authors:** Yoshitaka Uchida, Kawawa Banda, Imasiku Nyambe, Toru Hamamoto, Yui Yoshii, Kabenuka Munthali, Mukuka Mwansa, Moses Mukuka, Mubanga Mutale, John Yabe, Haruya Toyomaki, Yohannes Yared Beyene, Shouta M. M. Nakayama, Nobuyasu Naruse, Mayumi Ishizuka, Yukihiro Takahashi

## Abstract

Heavy metal contamination is a serious issue in many post-mining regions around the world. Kabwe town, Zambia, is known as one of the most polluted cities in the world, where high lead (Pb) levels have been reported in soils, plants, animals and human blood. Multidisciplinary approaches are critically needed to understand the current situation and to remediate the polluted area. In the current research, a large-scale preliminary field survey was performed to understand the current situation in Kabwe and to plan future mitigation approaches. Three aspects were mainly observed; 1) plant communities during the dry season in Kabwe city, 2) spectral images of the land surfaces in various locations in Kabwe and 3) Pb concentrations in soils and water. Overall, >15 different plant species were observed and many of them maintained their green colour even during the dry season. Some tree species, for example, *Caesalpiniaceae* and *Fabaceae* families may be utilised as phytostabilization approaches although their impacts on the soil Pb mobility should be further studied. For the spectral images, we used a handmade portable spectrometer, and our obtained spectral images showed typical curves observed from soils. These data may be used to understand the distribution of different soil types in this area, using aboveground images such as satellite images. For Pb concentrations in soils, extremely high total Pb levels (>1,000 ppm) was observed only within 2 km from the mining site. There was a weak but a positive correlation between the total and soluble Pb thus further study should also focus on the mobility of Pb from soils to plant ecosystems.

## 1. Introduction

There have been many studies done to evaluate the status of heavy metal contamination in Kabwe, Zambia. Due to the history of mining of lead (Pb) and zinc (Zn), it was reported that soils were contaminated with not only by Pb and Zn but also withcopper (Cu) and cadmium (Cd) (Nwankwo & Elinder, 1979; Tembo, Sichilongo, & Cernak, 2006). The contamination of the soils in the area resulted in extensive heavy metal pollution of livestock (Yabe et al., 2011; Yabe et al., 2013), vegetables, and humans (Clark, 1977; Yabe et al., 2015).

Some remediation approaches are needed in this area. Although the number of scientific reports is small, it has been identified that a phytoremediation approach could be used in Kabwe, where plants are used to reduce the risks related to the soil contamination (Reilly & Reilly, 1973; Leteinturier et al. 2001). Plants can remediate heavy metals using different mechanisms, for example, they may be able to extract heavy metals from the soil, and the plants can be taken away from the area to cleanse the contaminated soils. Contrastingly, plants are used to stabilize the heavy metals in soils, preventing them from escaping to other ecosystems (e.g. to groundwater or other soil surfaces as dust). Either way, the societal implementation of the phytoremediation approaches can be efficient when locally available or indigenous species are used.

Thus, a better understanding of the plant community structures for phytoremediation is needed in Kabwe. Earlier intensive surveys that were conducted for plant communities around the mining area in Kabwe recorded nearly 40 species (Leteinturier et al. 2001). However, these previous surveys were performed within the mining site, not extending to the whole Kabwe township. Also, the survey was performed at the end and the beginning of the wet season (April and November), thus not providing information for the driest period of the year (i.e. July–October). We believe that plants surviving in the dry season are critically important regarding their potential use as phytoremediation methods. Plants that become dominant during the dry season might be difficult to efficiently stabilise the Pb-containing soils because it was observed that extensive amount of fine soil particle (dust) is blown off during the dry season and plants may reduce the amount of dust.

Additionally, the adaptation (survival) of plant species and their efficacy regarding soil stabilisation are often influenced by soil types. Thus, detailed soil mapping is very important. Spectral sensing approaches can be an option to differentiate soil types according to their spectral curves and are often used using satellite data (Nanni et al. 2012). Soil properties such as soil moisture contents can also be quantified using the spectral data (Weidong et al. 2002). Also, the availability (mobility) of Pb in soils is another factor that must be considered to plan efficient phytoremediation approaches. Generally, in soils, Pb that is available to plants is very small when compared to the total Pb in soil. Thus, a better understanding of the amount of available Pb in soils is required not only to investigate the plant survival rates but also to investigate the risk of Pb leaching to water ecosystems (Gleyzes et al. 2002).

Thus, in the present study, plant community structures were investigated as well as land spectral images at various places in Kabwe town. Moreover, some basic information on soil Pb characteristics were studied.

## 2. Research methods

### 2.1 Research sites

The field survey was conducted from 5^th^ July to 8^th^ July 2016, in Kabwe town, Zambia. The sites where soil and plant samples were collected are shwn in Fig. 1. From the mine residue damping zone (“022_DUMP”, 14°27′44′′S, 28°25′51′′E), 0–5 km zones were covered in all the directions. The detailed description of each site is also stated in Table 1. Due to scarce rainfall during this period, most of the sites were dry, but some wetlands were found, particularly in northern part of the town (sites “015_dambo”, “016_dambo”, and “017_dambo_bridge”), where soils were saturated (Photo 1).

**Figure 1.**
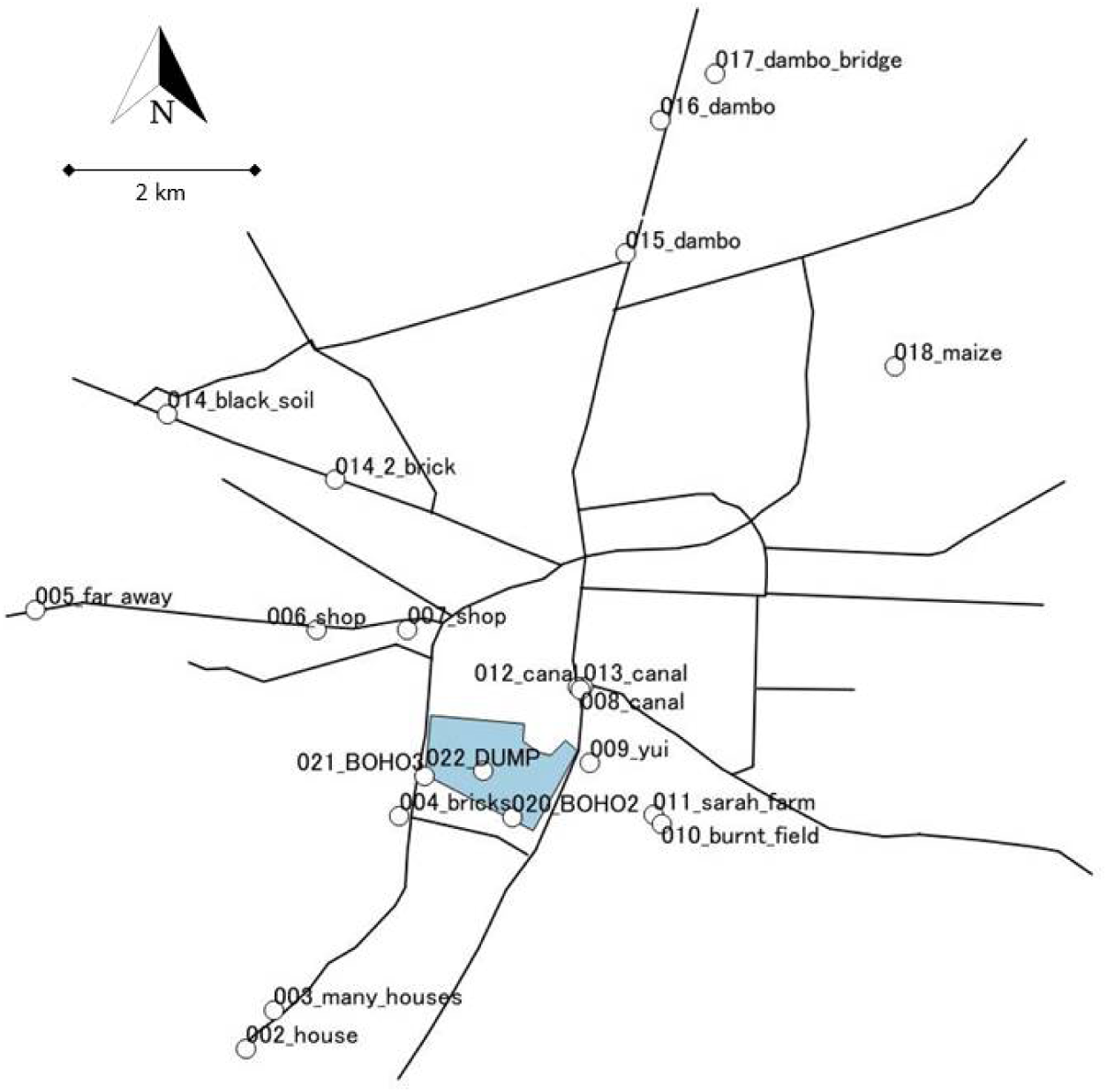
Major roads in Kabwe town, Zambia and our field survey sites (white circles). The shaded area was the mine residue dump site. The centre of the sampling sites, “022_DUMP” was at 14°27′44′′S, 28°25′51′′E. The map is North on the top, and the 2 km scale bar is indicated.

**Table 1.** Detailed description of the field survey sites.

| Number | Site name | Plant community information | Spectral image data | Soil XRF data | Soluble Pb data | Distance from the mine (m) |
| --- | --- | --- | --- | --- | --- | --- |
| 1 | 002_house | No | Yes | Yes | Yes | 3,787 |
| 2 | 003_many_houses | No | Yes | Yes | Yes | 3,298 |
| 3 | 004_bricks | No | No | Yes | Yes | 987 |
| 4 | 005_far_away | No | Yes | Yes | Yes | 4,956 |
| 5 | 006_shop | Yes | No | No | No | 2,275 |
| 6 | 007_shop | No | No | No | No | 1,674 |
| 7 | 008_canal | No | No | Yes | Yes | 1,334 |
| 8 | 009_yui | No | No | Yes | Yes | 1,112 |
| 9 | 010_burnt_field | Yes | No | Yes | Yes | 1,928 |
| 10 | 011_sarah_farm | No | No | Yes | Yes | 1,821 |
| 11 | 012_canal | No | No | No | No | 1,323 |
| 12 | 013_canal | No | No | Yes | Yes | 1,349 |
| 13 | 014_black_soil | Yes | Yes | Yes | Yes | 4,954 |
| 14 | 014_2_brick | Yes | Yes | No | No | 3,391 |
| 15 | 015_dambo | No | No | No | Yes | 5,556 |
| 16 | 016_dambo | Yes | No | No | No | 6,903 |
| 17 | 017_dambo_bridge | Yes | No | No | No | 7,626 |
| 18 | 018_maize | Yes | Yes | Yes | Yes | 5,993 |
| 19 | 020_BOHO2 | Yes | No | Yes | No | 577 |
| 20 | 021_BOHO3 | No | No | Yes | No | 613 |
| 21 | 022_DUMP | Yes | No | Yes | No | 0 |

**Photo 1.**
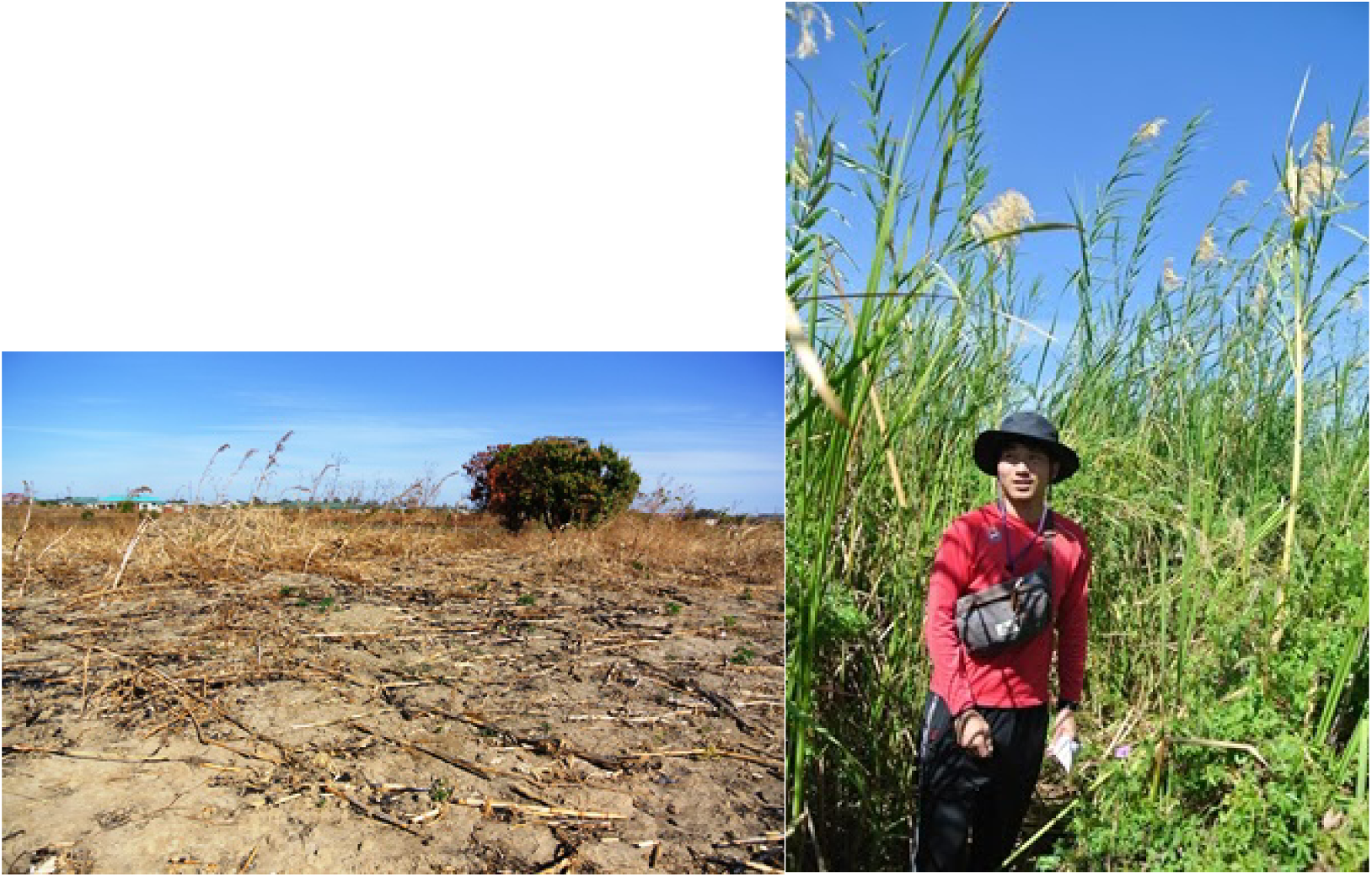
(Left) Dry field (site “018_maize”) and (Right) wetland site (site “016_dambo”) (taken on July 6^th^, 2016).

### 2.2 Plant survey

At each plant survey site (Fig. 1), plant species were visually observed. To identify species, van Wyk and van Wyk (2013) and Pienaar and Smith (2011) were referred to as well as communications with locals.

### 2.3 Measurement of soil Pb concentrations using X-ray fluorescence approaches (XRF) and a portable colorimetric Pb measurement kits

Approximately 200 g of soils were collected from each soil sampling site. They were placed in plastic bags (Ziploc, Asahi Kasei Home Products Corporation, Japan) and measured for total Pb concentrations using a portable XRF analyser (Innov-X Alpha Series™ Advantages, Innov-X Systems, Inc. USA). Then, the metals in soils were extracted with hot water (80°C, 1 hour, soil:water = 1:10). The supernatant was analysed for the soluble Pb concentrations using a colorimetric kit. Also, some surface and groundwater were analysed for soluble Pb, using the same kit.

### 2.4 Spectral sensing

A portable spectral sensor was used to investigate the spectrum of land surfaces in different areas in Kabwe. The compact hyperspectral sensor with a USB camera (MCM-4304, Micro Vision Co., Ltd. Japan) was used. The measurement was taken using following approaches: first, a total reflection at each soil measurement site was obtained by taking a picture of a white filter paper (Tokyo Roshi Kaisha, Ltd. Japan). Then, a soil reflectance was measured without moving the sensor (the sensor was secured on a stand, as shown in Photo 2). Five replicates were taken for each sample.

**Photo 2.**
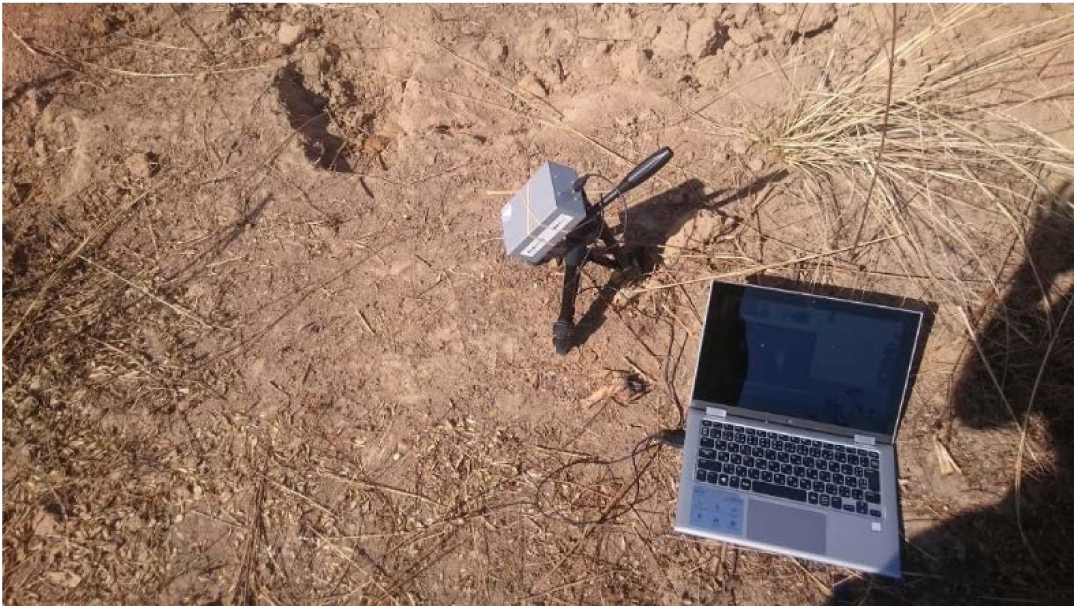
The compact hyperspectral sensor with a USB camera. The camera was set up at a right angle from the sun to avoid the shadow.

For data analyses, ImageJ software (ver. 1.6.0_24, National Institutes of Health, USA) was used. First, the spectral data was converted to values depending on the strength of white. Then, the X-axis was adjusted for the wavelength using a Hg light value. Spectral data of Hg light has some specific peaks, and these peaks are known as specific wavelengths (Sansonetti et al. 1996). To show the data as reflectance, the data from the soil samples were divided by the spectral data taken from a white filter paper.

## 3. Research activities and snapshots of the preliminary findings from the field survey in Kabwe

### 3.1 Observed plant species

The plant communities in Kabwe were recorded in detail by Leteinturier et al. (2001). Most of the plants found in the present study were also reported in the previous report. As shown in Table 1 a total of 20 plant species were identified in the present study. The survey conducted in the current study was not intensive enough to discuss the effect of the mine residue on plant community structures or diversities. On the mine residue, many African fountain grass species (*Pennisetum setaceum*) was observed. According to verbal communication with the landowners, this species has been markedly increasing over the last decades and helped to reduce the dust blowing around the residues. In a wetland site (site 016), totally different plant community structures were observed, including *Phragmites* and *Ipomoea*. This suggests that the availability of water is probably one of the major factors controlling the plant community structures in Kabwe area. Further studies must focus on plant characteristics regarding the mobility of heavy metals.

**Table 1.** Plant species observed in Kabwe, Zambia, during the dry season (July 2015). Plant species were visually determined according to Leteinturier et al. (2001), van Wyk and van Wyk (2013) and Pienaar and Smith (2011). The “+” sign means that species were observed at each site. The site names were in the order of distance from the mine (m), from left to right.

|  | Distance from the mine<br>(m) | 0 | 613 | 1,112 | 1,349 | 2,275 | 3,391 | 4,954 | 5,556 | 5,993 | 6,093 |
| --- | --- | --- | --- | --- | --- | --- | --- | --- | --- | --- | --- |
| Species | Family/Site name | 022 | 020 | 009 | 013 | 005 | 014_2 | 014 | 015 | 018 | 016 |
| *Ageratum conyzoides* L. | Asteraceae |  |  |  |  |  |  |  | + |  |  |
| *Amaranthus dubius* Mart. ex Thell. | Amaranthaceae |  |  |  |  |  |  |  |  | + |  |
| *Argemone mexicana* | Papavaceae |  |  | + |  |  |  |  |  |  |  |
| *Bauhinia petersiana* bolle | Caesalpiniaceae |  |  | + | + | + | + |  |  |  |  |
| *Bidens oligoflora* (Klatt) Wild | Asteraceae |  |  |  |  |  | + |  | + |  |  |
| *Bidens pilosa* | Asteraceae |  |  |  |  |  |  | + |  |  |  |
| *Celosia trigyna* | Amaranthaceae | + |  |  |  |  |  |  |  |  |  |
| *Crotalaria agatiflora* | Fabaceae |  |  |  |  |  |  |  |  |  |  |
| *Cynodon dactylon* (L.) Pers. | Poaceae | + |  |  |  |  |  |  |  |  |  |
| *Cyperus involucratus* Rottb. | Cyperaceae |  |  |  | + |  |  |  |  |  |  |
| *Flaveria trinerva* (Spreng.) Mohr | Asteraceae | + |  |  |  |  |  |  |  |  |  |
| *Ipomoea cairica* (L.) Sweet | Convolvulaceae |  |  |  |  |  |  |  |  |  | + |
| *Lebeckia cytisoides* | Fabaceae |  |  |  | + |  |  |  |  |  |  |
| *Pennisetum setaceum* (Forssk.) Chiov. | Poaceae | + | + |  |  |  |  |  |  |  |  |
| *Phragmites mauritianus* L. | Poaceae |  | + |  |  |  |  |  |  |  | + |
| *Ricinus communis* var. *communis* | Euphorbiaceae |  |  |  |  |  |  | + |  |  |  |
| *Solanum incarnum* L. | Solanaceae |  |  |  |  |  |  | + |  |  |  |
| *Trichodesma zeylanicum* (Burm f.) R Br. | Boraginaceae |  |  |  |  |  |  | + |  |  |  |
| *Tridax procumbens* L. | Asteraceae |  |  |  |  |  |  | + |  | + |  |
| *Vernonia erinacea* Wild | Asteraceae |  |  |  | + |  |  | + |  |  |  |

### 3.2 Heavy metal concentrations in surface soils

Based on XRF measurements in the current study, the total Pb contents in soils ranged from 20 ppm (site 014_2_brick) to 10,000 ppm (site 008_canal). There was a weak but positive relationship between the total and water soluble Pb contents (Fig. 1a). The water soluble Pb contents ranged from 0 to 4 ppm and it was, in general, less than 1% of the total Pb. We believe that soil characteristics (e.g. parent materials and clay contents) are the major factors controlling the ratio between the total and water soluble Pb. Further research is needed in this area because water soluble Pb can be more mobile and pose a higher risk to plant and animal absorption.

**Figure 2.**
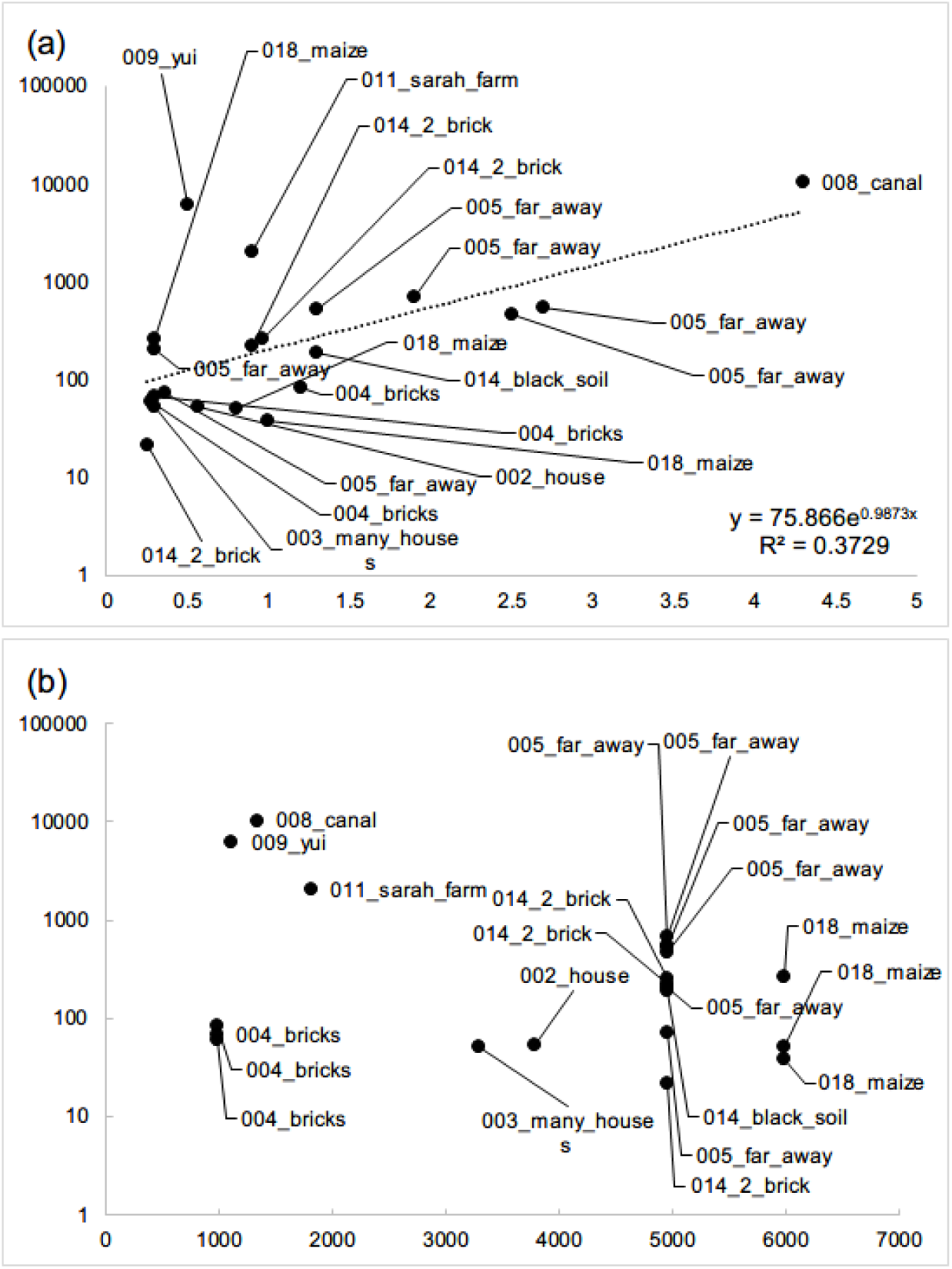
(a) The correlation between water soluble Pb concentration and total Pb concentration and (b) the correlation between the total Pb concentration and the distance from the mine site (site 022). The Y axis were shown as a log-scale. For a detailed description of the site, refer to Fig. 1 and Table 1.

### 3.3 Spectral characteristics of the site

Fig. 3 shows photos of some soil samples with their reflectance data. The soil from site 005 was brighter in colour and its spectral data shows a marked difference compared with other soils. Moreover, soil samples from site 014 showed a different pattern. Based on personal communications, local people valued this soil as a relatively more fertile soil (called “black soil”) and the identification of this soil, when compared with other soils, seemed promising based on preliminary data from the current study. The data below 400 nm or above 900 nm showed large variabilities, but the data between these values (i.e. 400–900 nm) could be used to differentiate the soils. The influence of soil moisture was ignored since the soils were very dry but future analyses should also focus on the effects of soil moisture.

**Figure 3.**
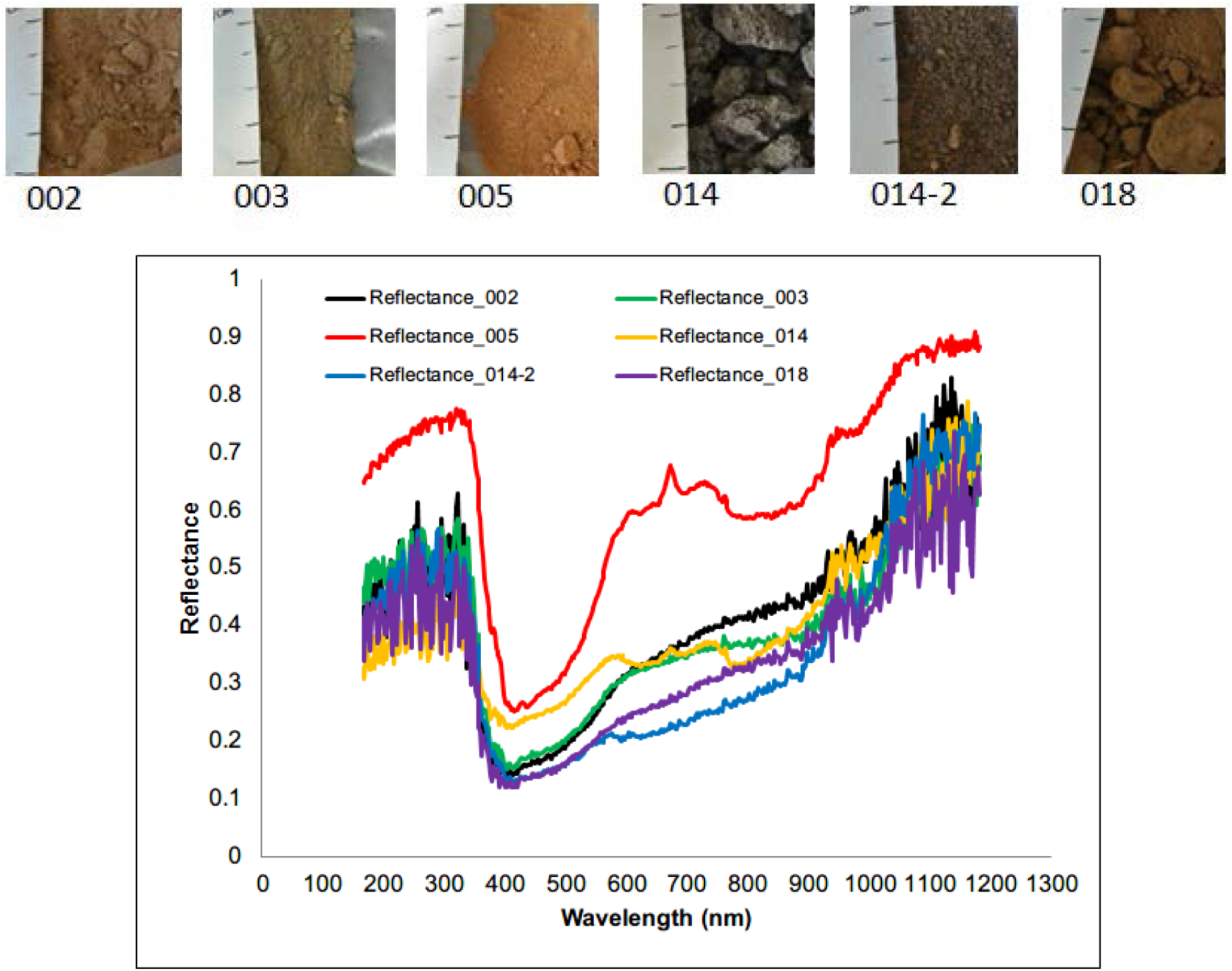
Photos of some of the sampled soils in Kabwe (top) and their reflectance (bottom).

## 4. Conclusion

The preliminary field survey in the current study clearly indicated that an extensive area within Kabwe was contaminated with Pb. Plant community structure observation revealed that plant species such as *Caesalpiniaceae* and *Fabaceae* families remained viable during the dry season and could potentially be utilised to stabilise the contaminated soils. The percentage of water soluble Pb in the total soil Pb was < 1 %, but the extreme values were observed within 2 km zone from the mining site. The preliminary taken spectral curves suggested that remote-sensing techniques may be used to differentiate soil types. Further studies should focus on more detailed analyses of plant communities as well as their capacities to stabilise soils and Pb. Also, soil moisture has to be taken into account to further evaluate the potential use of spectral images in the area.

## Acknowledgements

We acknowledge the support from Russell Dowling (Pure Earth) and Jack Caravanos (Pure Earth) for XRF measurement. This research was supported by JST/JICA, SATREPS (Science and Technology Research Partnership for Sustainable Development).

